# A Draft Pacific Ancestry Pangenome Reference

**DOI:** 10.1101/2024.08.07.606392

**Authors:** Connor Littlefield, Jose M. Lazaro-Guevara, Devorah Stucki, Michael Lansford, Melissa H. Pezzolesi, Emma J. Taylor, Etoni-Ma’asi C. Wolfgramm, Jacob Taloa, Kime Lao, C. Dave C. Dumaguit, Perry G. Ridge, Justina P. Tavana, William L. Holland, Kalani L. Raphael, Marcus G. Pezzolesi

## Abstract

Individuals of Pacific ancestry suffer some of the highest rates of health disparities yet remain vastly underrepresented in genomic research, including currently available linear and pangenome references. To begin addressing this, we developed the first Pacific ancestry pangenome reference using 23 individuals with diverse Pacific ancestry. We assembled 46 haploid genomes from these 23 individuals, resulting in highly accurate and contiguous genome assemblies with an average quality value of 55.0 and an average N50 of 40.7 Mb, marking the first *de novo* assembly of highly accurate Pacific ancestry genomes. We combined these assemblies to create a pangenome reference, which added 30.6 Mb of novel sequence missing from the Human Pangenome Reference Consortium (HPRC) reference. Mapping short reads to this pangenome reduced variant call errors and yielded more true-positive variants compared to the HPRC and T2T-CHM13 references. This Pacific ancestry pangenome reference serves as a resource to enhance genetic analyses for this underserved population.

## Introduction

The human reference genome has been foundational in our understanding of human genetic variation since the publication of the first human genome sequence in 2001^1^. Efforts to refine this reference have led to significant advancements, including the recent completion of the first complete haploid genome reference, Telomere-to-Telomere CHM13 (T2T-CHM13)^2^. These developments have greatly enhanced our ability to study genetic variation on a genome-wide scale. Nonetheless, a single reference genome cannot fully capture the genetic diversity present in human populations worldwide. As a result, mapping sequence data to these references introduces reference bias and variant calling errors, particularly in genomic regions with structural variations or high levels of polymorphism^3–5^. These issues are especially pronounced in sequence data from individuals belonging to populations not represented in these reference genomes.

To address the limitations of single-reference genomes, initiatives like the Human Pangenome Research Consortium (HPRC) are developing pangenome references that incorporate many genomes from individuals of diverse ancestries^6^. New advancements in long-read DNA sequencing technologies and the development of sophisticated graph-formatted references have made this possible, significantly reducing the time and resources necessary to achieve it^7–10^. Pangenome references offer a more accurate representation of complex structural variation and enhance short-read mapping and variant identification, particularly in medically relevant and traditionally challenging genomic regions^6,11,12^. This allows for downstream disease association and functional analyses of previously inaccessible variants, many of which overlap with clinically significant genes that may contribute to disease. Additionally, the recent development of population-specific pangenomes, such as the draft Chinese^12^ and Arab^11^ pangenome references, has enhanced the analysis of genetic variation unique to populations. These pangenomes include novel sequences and variants not represented by the HPRC pangenome graph alone, improving the discovery of variants relevant to population-specific disease studies.

Individuals of Pacific ancestry have been underrepresented in genetic research, with this population excluded from existing pangenome references and extensive sequencing projects such as the 1000 Genomes Project^13^. To date, less than 0.1% of genome-wide association studies (GWAS) have included individuals of Pacific ancestry^14^. Despite this, individuals of Pacific ancestry experience disproportionately high rates of metabolic diseases, such as obesity^15^, Type 2 diabetes (T2D)^16,17^, and kidney disease^18^, with a suspected genetic component contributing to these elevated rates^19^. The scarcity of genetic cohorts of Pacific ancestry, especially those with whole-genome sequence data, has been the main barrier to advancing genetic research in this population. In response to this gap, we and others are increasing sequencing efforts in this population to better understand the genetic basis of metabolic disease^20–22^. Here, we aimed to create a Pacific ancestry pangenome reference as a valuable community resource to enhance variant calling and offer a comprehensive examination of genetic variation in this underrepresented population.

## Results

### Assembling 46 Pacific Ancestry Genomes

Twenty-three participants from the Utah Pacific Islander Diabetes Study were included in this study. These participants had diverse Pacific ancestry—Tonga (6), Samoa (6), Fiji (4), the Philippines (2), the Marshall Islands (2), Guam (1), Tahiti (1), and Pohnpei (1)—and reported that all four of their grandparents originated from the same Pacific Island (Fig. 1a, Supplementary Table 1). All participants provided informed written consent, and the study was approved by the University of Utah Institutional Review Board (IRB_00159221).

**Figure 1.**
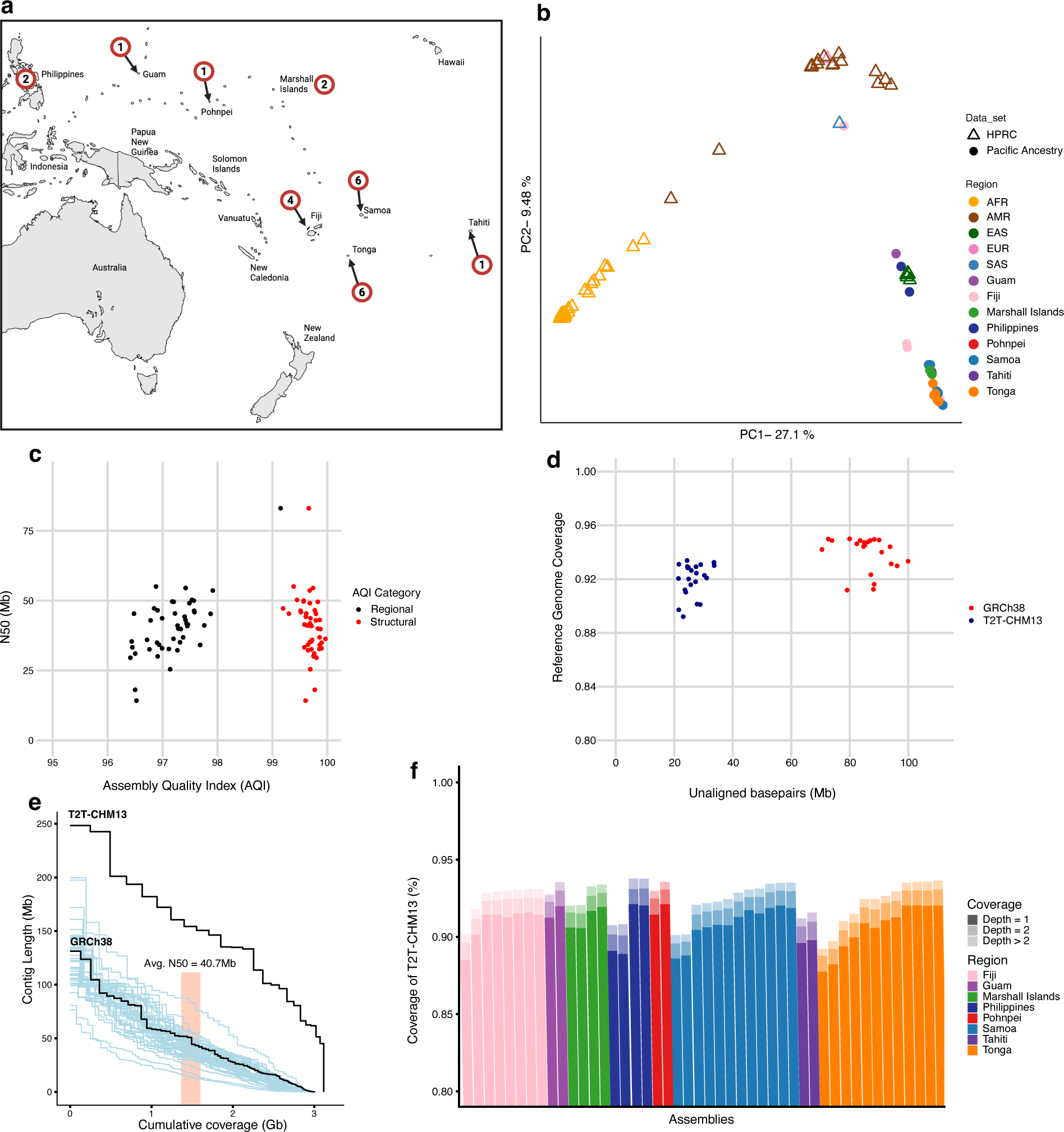
Assessing the quality of 46 partially-phased haploid Pacific ancestry assemblies. **a)** Map showing the reported ancestry of the 23 individuals selected for long-read sequencing and de novo genome assembly. Each of these participants reported that all four grandparents originated from the same region. The numbers in red circles indicate the number of individuals with ancestry from each region. **b)** Principal component plot of variation in the 47 HPRC assemblies and the 23 Pacific ancestry assemblies. **c)** Quality assessment of the 46 Pacific ancestry assemblies. *X* axis shows the CRAQ Assembly Quality Indicator (AQI) values for structural quality (red) and regional quality (black), while the *Y* axis shows the N50 values for each assembly. **d)** Reference-based quality assessment. The *X* axis shows the number of base pairs from each haploid assembly that do not align to linear references GRCh38 (red) and T2T-CHM13 (blue). The *Y* axis shows the percentage of the reference genome covered by at least one alignment for each assembly. **e)** Assessing contiguity. The black lines represent the linear references T2T-CHM13 and GRCh38 (N-Split), while the blue lines represent single Pacific ancestry haploid assemblies. Each step indicates a single contig in the assembly. The *X* axis shows the cumulative length of the assembly contigs and the *Y* axis shows the individual contig length. **f)** Individual assembly coverage of T2T-CHM13. The *Y* axis shows the percentage of T2T-CHM13 covered for each Pacific ancestry assembly on the *X* axis.

Whole-genome sequencing was performed on DNA from whole-blood for each participant, generating PacBio high-fidelity (HiFi) long reads at an average depth of 32.57x (range: 24.00x-56.55x) and highly accurate Illumina short reads at an average depth of 32.74x (range: 30-38x). Principal component analysis of the genetic variation within these samples demonstrates the distinct diversity they contribute to the existing Human Pangenome Reference Consortium (HPRC) assemblies^6^ (Fig. 1b).

We developed a genome assembly pipeline incorporating Hifiasm^8^ to create 23 primary assemblies and 46 partially-phased haploid assemblies using the HiFi long reads. These assemblies were subsequently evaluated and refined using Inspector^7^, which aligns HiFi long reads from each sample to its respective assembly to calculate quality metrics, identify errors, and correct them through local reassembly. The quality of the corrected 46 haploid assemblies was assessed by evaluating the contiguity, completeness, and accuracy of each assembly. Contiguity was measured using N50, the length of the shortest contig at 50% of the total assembly length. The average N50 value for the 46 assemblies was 40.7 Mb (range: 14.2 Mb– 83.0 Mb), comparable to the linear human reference GRCh38 (58 Mb) (Fig. 1c,e). Completeness was assessed by the total length of each assembly, which averaged 2.98 Gb (range: 2.86 Gb–3.03 Gb) (Supplementary Table 1). This corresponds to 95.5% of the length of the most complete linear human reference, T2T-CHM13, which is 3.12 Gb. Completeness was further evaluated by examining the percentage of expected protein-coding genes in each assembly via compleasm^23^. Using this method, assembly completeness ranged from 92.1% to 99.2% with an average of 96.6% (Supplementary Fig. 1, Supplementary Table 2).

The accuracy of the assemblies was assessed using the HiFi reads mapped to their respective assemblies by Inspector. For each of the 23 samples, 100% of the HiFi reads remapped to the primary assembly, and on average, 99.93% of these reads remapped to each respective haploid assembly. On average, the assemblies exhibited 10.7 (range: 2–22) structural errors (>50bp) and 4,859.1 (range: 3,009–10,744) small errors (<50bp) (Supplementary Fig. 2a,b). The Phred-scaled quality values (QV) for the assemblies averaged 55.0 (range: 51.1–58.6), which is approximately one error every 300,000bp (Supplementary Fig. 3b). To avoid potential bias from using Inspector both for correcting and assessing assembly errors, we also employed CRAQ^10^ to further, and independently, assess assembly quality. Unlike Inspector, which relies solely on long reads, CRAQ utilizes both highly accurate short reads and HiFi long reads mapped to their assemblies to identify errors. CRAQ detects errors by analyzing clipped reads to identify regional and structural issues, and it provides an overall score called the Assembly Quality Indicator (AQI). This method helps reduce the effect of error-prone regions on the overall quality score and distinguishes between smaller regional errors and larger structural misjoins. The average regional AQI (R-AQI) for the 46 haploid assemblies was 97.19 (range: 96.41–99.15), and the average structural AQI (S-AQI) was 99.69 (range: 99.20–99.97) (Fig. 1c). For reference, the Genome in a Bottle (GIAB) HG002 sample assembled using HiFi reads (∼55x) and Canu showed an R-AQI of 98.96, an S-AQI of 93.92, and a QV of 46.57^10^.

Both Inspector and CRAQ are reference-free tools designed to evaluate assembly accuracy. However, to assess assembly accuracy, we can also map assembly contigs to high-quality linear reference genomes. Although individuals of Pacific ancestry might have unique sequences that do not align with current reference genomes, we would still expect most of the sequences in a high-quality assembly to align with these references. Mapping the contigs of our 46 haploid assemblies to both GRCh38 and T2T-CHM13 revealed that, on average, 93.50% (Supplementary Fig. 4) and 92.40% of these genomes, respectively, were covered by at least one alignment. This indicates that portions of the Pacific ancestry assemblies are systematically unassembled. For T2T-CHM13, 90.82% of the genome was covered at a depth of 1, and 1.58% was covered at a depth ≥ 2 (Fig 1f), representing duplicated regions in the Pacific ancestry assemblies. Additionally, by examining the portions of the Pacific ancestry assemblies that did not map to the references, we can identify novel sequences within our assemblies. An average of 84.00 Mb (range: 66.92 Mb–103.96 Mb) did not align to GRCh38, and 26.87 Mb (range: 18.73 Mb–38.81 Mb) did not align to T2T-CHM13 (Fig. 1d). Of the sequences unaligned to T2T-CHM13, an average of 77.59% were annotated by RepeatMasker^24^ as repetitive sequences, with 43.48% being satellite DNA. These unaligned sequences represent putative novel sequences unique to individuals of Pacific ancestry.

### Copy Number Variation in the 46 Assemblies

We leveraged these highly accurate assemblies to examine copy number variants (CNVs) in our 23 Pacific ancestry genomes. To do this, we used Liftoff^25^ to annotate protein-coding genes in each haploid assembly and identified those genes that were marked as having at least one extra copy. We performed the same analysis on the 94 assemblies published by the HPRC and compared these results to our Pacific ancestry assemblies to identify population-specific differences in protein-coding gene duplications. In the Pacific ancestry assemblies, we identified 50–100 duplicated genes per individual, totaling 704 unique whole-gene duplications (Fig 2e). Among these, 326 were specific to individuals of Pacific ancestry (Fig 2d), with 39 found in two or more assemblies. Further examination of the 39 duplications unique to Pacific ancestry assemblies revealed that three assemblies had duplications of the *NPHP1* gene, which is associated with renal disease^26^ (Fig. 2c). We next observed differences in gene duplication frequencies between Pacific ancestry and HPRC assemblies. 29 genes had more than a 10% frequency increase in Pacific ancestry assemblies (Fig. 2a), while 56 genes showed a similar increase in HPRC assemblies (Fig. 2b). Alternatively, we examined duplications that were more frequent in the HPRC assemblies and found that the *CLPS* gene, which encodes pancreatic colipase and is linked to Type 2 Diabetes^27^, was duplicated in 57% of HPRC assemblies but only 32% of Pacific ancestry assemblies. These Pacific ancestry-specific patterns of duplicated genes linked to metabolic diseases highlight the potential role of copy number variation in modifying metabolic disease susceptibility in this population.

**Figure 2.**
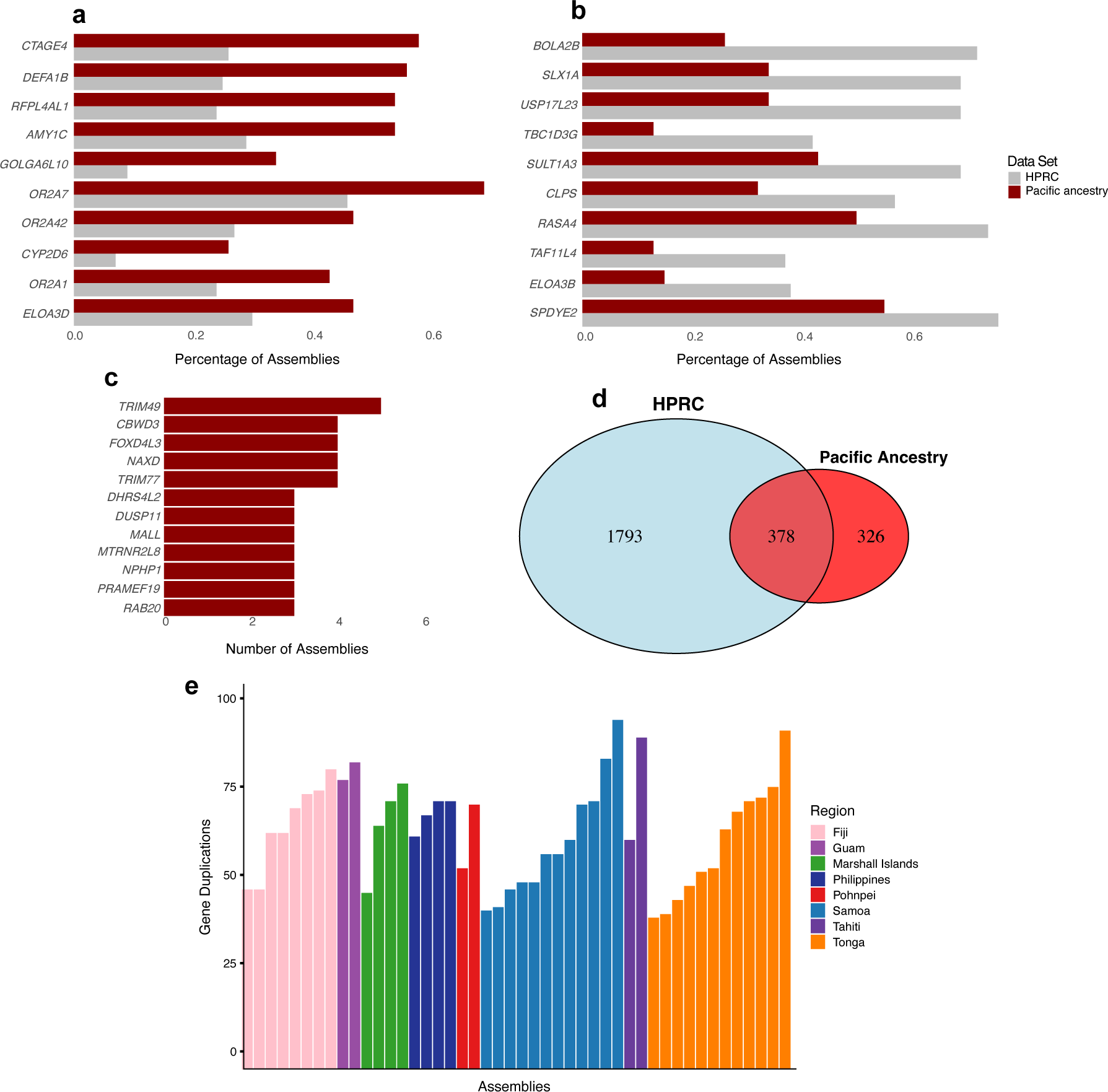
Comparing gene duplications in the 94 HPRC and 46 Pacific ancestry assemblies. **a)** The top ten protein-coding genes duplicated more frequently in Pacific ancestry assemblies compared to HPRC assemblies. **b)** The top ten genes duplicated more frequently in HPRC assemblies. **c)** Gene duplications present only in the Pacific ancestry assemblies and absent from all HPRC assemblies. **d)** Venn diagram showing the number of genes duplicated only in the HPRC or Pacific ancestry assemblies, and those shared between the two. **e)** The number of genes duplicated per Pacific ancestry assembly, ordered by Pacific ancestry

### A Pacific Ancestry Pangenome Graph Reference

To overcome reference bias inherent in the linear reference genomes and improve variant calling in individuals of Pacific ancestry, we utilized the 46 Pacific ancestry assemblies to create a pangenome graph reference using the Minigraph-Cactus pipeline^28^ (Fig 3a, Supplementary Fig. 7). Minigraph-Cactus generates assembly-based pangenome graphs by aligning de novo assemblies to a reference. The process begins by utilizing Minigraph to create a structural-variation-only graph. The assemblies are then remapped to this graph to incorporate single nucleotide variations, while sequences longer than 10 kb that remain unaligned are excluded to remove highly repetitive regions, such as those in telomeres and centromeres. We applied this pipeline to our 46 haploid assemblies by mapping each assembly and GRCh38 to the linear T2T-CHM13 reference. The resulting pangenome graph comprises 47 million nodes and 65 million edges and has a complexity value of 1.37 (nodes/edges). The total base pair length of the graph is 3.22 Gb, which is the base pair sum of all nodes. We found that the 46 Pacific ancestry assemblies added 92.5 Mb of novel sequence to the GRCh38 and T2T-CHM13 references, with 60.6 Mb of this present in more than one sample (Fig. 3b,c). Notably, 1.4 Mb of this sequence was seen in 95% of samples, representing the core Pacific ancestry sequence missing from GRCh38 and T2T-CHM13.

**Figure 3.**
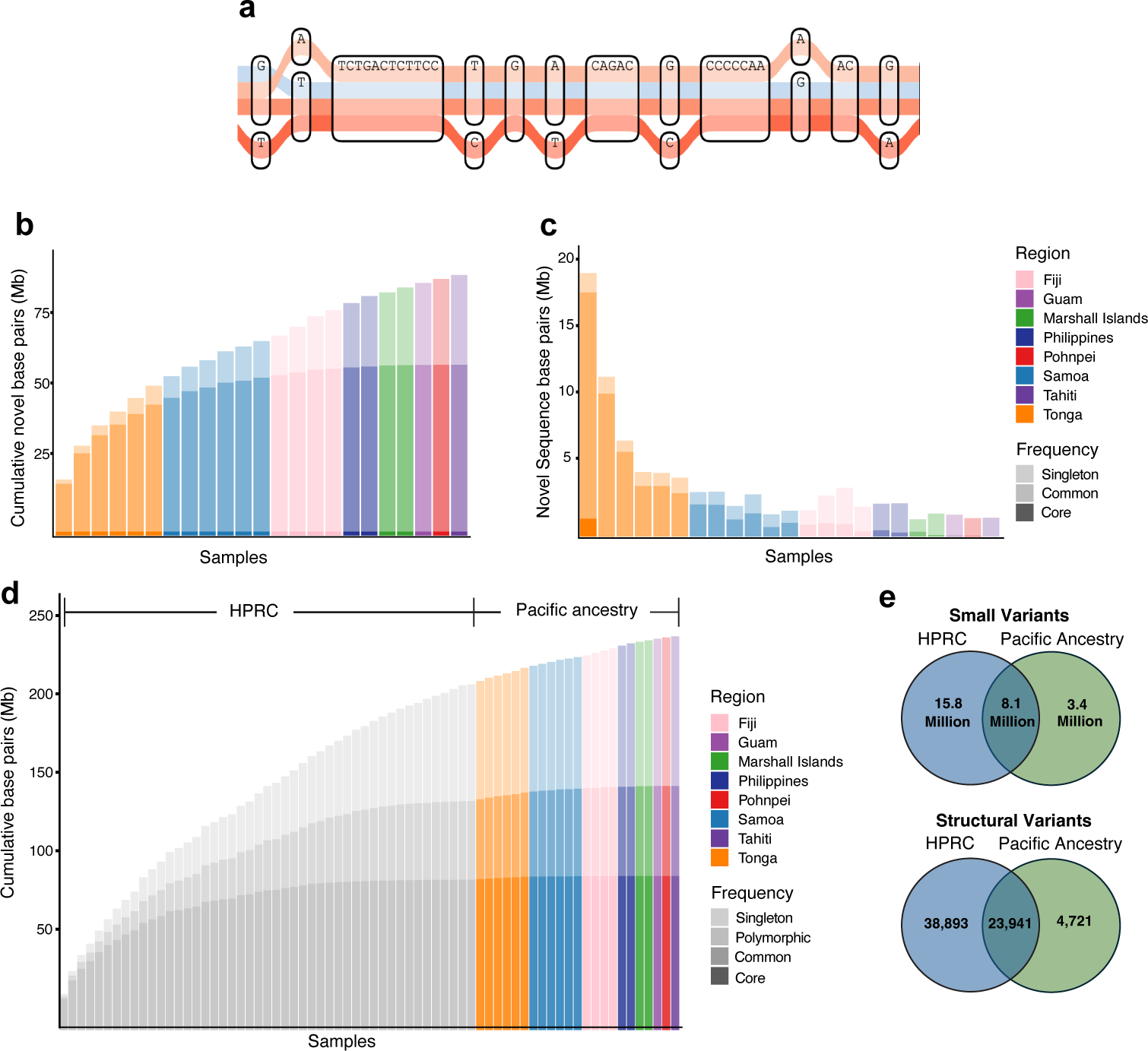
Examining Pacific ancestry-specific variation in the pangenome graphs. **a)** Visual representation of four representative haplotypes in the Pacific ancestry pangenome variation graph at the HLA-A region. The DNA sequence boxes represent nodes, and the colored lines passing through each node indicate individual haplotypes. The red colored paths indicate Pacific ancestry haplotypes, while the blue path represents the GRCh38 reference. **b)** Cumulative number of novel base pairs added to T2T-CHM13 and GRCh38 as each Pacific ancestry sample is integrated into the pangenome graph. The color intensity indicates the frequency of the novel sequence: singleton sequence (observed only once), common sequence (observed in at least 5% of samples), and core sequence (observed in at least 95% of samples). **c)** Number of novel base pairs contributed by each individual assembly to the pangenome variation graph. **d)** Cumulative novel base pairs added to T2T-CHM13 and GRCh38 references by HPRC samples (gray) and Pacific ancestry samples (colored), ordered by ancestry in the combined HPRC + Pacific ancestry variation graph. Polymorphic sequence refers to those sequences observed more than once in the variation graph. Core sequence in this graph is very low (0.3Mb) and not visible. **e)** Variants decomposed from the combined HRPC + Pacific ancestry graph stratified by size into structural (>50bp) and small (<50bp) variants. Variants in the green represent those observed only in the Pacific ancestry assemblies.

We next aimed to determine how much of the novel sequence in our Pacific ancestry pangenome reference was also missing from the HPRC reference. To do this, we created a combined Pacific ancestry + HPRC pangenome graph by aligning the 94 HPRC assemblies, 46 Pacific ancestry assemblies, and GRCh38 to the T2T-CHM13 reference. This combined graph contained 104 million nodes and 144 million edges, had a complexity value of 1.38, and a total base pair length of 3.38 Gb. Our analysis showed that the addition of Pacific ancestry samples contributed 30.6 Mb of novel sequence to the pangenome graph (Fig 3d, Supplementary Fig. 5). Of this, 9.6 Mb were observed at least twice among the samples, representing Pacific ancestry-specific polymorphic sequences. In this combined graph, the core sequence (95% of samples) dropped to 0.3 Mb, highlighting core genetic variation missing from GRCh38 and T2T-CHM13.

To better understand the variation contributed by the Pacific ancestry assemblies, we deconstructed the combined Pacific ancestry + HPRC graph to identify variants unique to individuals of Pacific ancestry. In total, we identified 26.1 million variants in the combined pangenome. Of these, 8.1 million variants were shared between the HPRC and Pacific ancestry assemblies, while 3.4 million variants were specific to the Pacific ancestry assemblies. We further stratified these Pacific ancestry-specific variants by size and identified 3,436,482 small variants (<50 bp) and 4,721 structural variants (>50 bp) (Fig. 3e).

### Applying the Pacific Ancestry Pangenome

We next sought to quantify the benefits of the Pacific ancestry pangenome reference by aligning whole-genome short reads and measuring the improvements in small variant calling. To do this, we first needed to identify the pangenome graph allele frequency filters that optimized variant calling accuracy for both the HPRC and Pacific ancestry pangenome graphs. Measuring the accuracy of variant calls requires comparing the calls against a truth set. Since there are no well-characterized truth sets of variants for individuals of Pacific ancestry, we created a pseudo-truth set by aligning our most accurate Pacific ancestry assembly, sample UP009 (N50 values of 83.0 Mb and 53.6 Mb for haplotypes 1 and 2), to T2T-CHM13 and calling variants via Dipcall^29^. We then aligned the Illumina whole-genome short-read sequencing reads for sample UP009 to the HPRC and Pacific ancestry references and called small variants (<50bp). We measured the concordance of the short read call set with the pseudo-truth set, limiting the analysis to confident regions identified by Dipcall. This process was repeated for multiple versions of the Pacific ancestry and HPRC graph pangenomes filtered by allele frequency. Our analysis revealed that an allele frequency filter of 2 yielded the highest F1 score for the Pacific ancestry reference, while an allele frequency filter of 4 resulted in the highest F1 score for the HPRC reference (Fig. 4d,e). We thus decided that further benchmarking would be limited to these two allele-frequency-filtered graphs.

**Figure 4.**
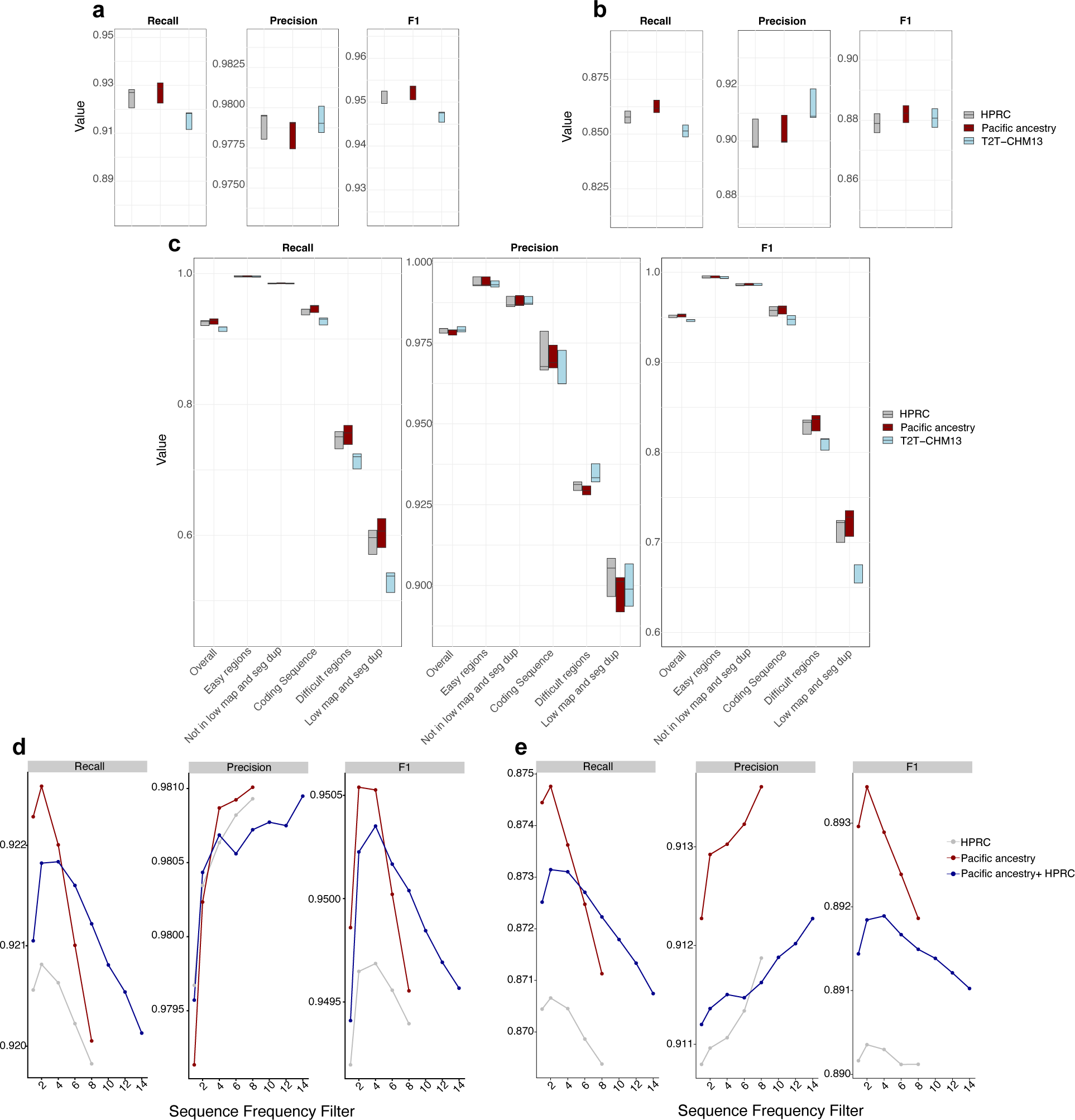
Evaluating improvements in small variant calling by aligning whole-genome short read sequences to the Pacific ancestry pangenome reference. **a)** Box plots showing recall, precision, and F1 scores for SNVs called from short read sequence data of five Pacific ancestry samples genome wide. The samples are aligned to either the HPRC pangenome reference (gray), the Pacific ancestry pangenome reference (red), or T2T-CHM13 (blue). **b)** Same box plots as in (a), but for indels instead of SNVs. **c)** Box plots of recall, precision, and F1 scores for SNVs using the same data as in a), but stratified across different genomic regions specified by the Genome in a Bottle consortium (GIAB)^30^.”low map” refers to low mappability, and “seg dup” refers to segmental duplications. **d)** Recall, precision, and F1 scores for SNVs called from sample UP009 short read sequence data. The sample was aligned to the HPRC pangenome reference (gray), the Pacific ancestry pangenome reference (red), and a combined HPRC + Pacific ancestry pangenome reference (blue) across multiple allele frequency filters ranging from 0-14. **e)** Same plots as in (d), but for indels instead of SNVs.

We also tested if the combined Pacific ancestry + HPRC pangenome graph could improve variant calling compared to the Pacific ancestry graph alone. However, we found that the F1 scores for the combined graphs were consistently lower (Fig. 4d,e). This may be due to limitations in the Giraffe personalized pangenome algorithms to accurately select representative haplotypes when additional haplotypes of differing ancestries are available.

Having identified the ideal allele frequency filters for the pangenome graphs, we next sought to quantify the improvements in small variant calling using short-read sequences from Pacific ancestry samples. We selected the top five samples from the 23 participants based on assembly contiguity (N50). Again, we created pseudo-truth sets for these samples using their respective assemblies mapped to T2T-CHM13. We then mapped the Illumina short reads of these samples to the Pacific ancestry reference, the linear T2T-CHM13 reference, and the HRPC reference, and called variants. We analyzed the concordance between these variants and the pseudo-truth sets for single nucleotide variants (SNVs) and small insertions and deletions (indels) separately and stratified the concordance metrics by various genomic regions^30^. Aligning to the Pacific ancestry pangenome reduced the total number of variant call errors (false negatives + false positives) (Supplementary Fig 9a,b, and Supplementary Fig. 10a,b) and improved the average F1 score for both SNVs and indels in all five samples (Fig 4a,b). Recall scores were typically highest across all genomic regions for the Pacific ancestry reference, while precision scores were typically highest for the T2T-CHM13 reference (Fig 4c, and Supplementary Fig 7). The benefits of aligning to the Pacific ancestry pangenome were most pronounced in traditionally difficult-to-map genomic regions. On average, the additional number of true positive SNVs gained in all regions compared to the HPRC graph and the T2T-CHM13 reference was 8,811 and 41,308, respectively (Supplementary Fig 9a, Supplementary Table 3); for indels, the gain was 4,003 and 10,092 respectively (Supplementary Fig 10a, Supplementary Table 4).

## Discussion

The goal of this study was to comprehensively characterize genetic variation in individuals with Pacific ancestry and create a pangenome resource to improve variant calling in this population. We *de novo* assembled 46 partially-phased haploid genomes from 23 individuals of diverse Pacific ancestry. With these assemblies, we built a Pacific ancestry pangenome reference that improves variant calling accuracy using short-read sequence data from this population, allowing the identification of variants that were previously undetectable using other references.

Individuals with Pacific ancestry suffer disproportionately from metabolic diseases^15–18^, and the genetic contributions to these conditions are poorly understood, partly due to the lack of genotype and whole-genome sequence data for this population. We and others are advancing efforts to better understand the genetic underpinnings of metabolic disease in individuals with Pacific ancestry^20–22^; this pangenome reference further enhances these efforts. It improves the applicability of short-read sequence data to accurately identify genetic variation in individuals with Pacific ancestry, providing a valuable resource for further genetic research in this population.

To build this resource, we sequenced the genomes of 23 individuals with Pacific ancestry, obtaining PacBio HiFi reads at an average depth of 32.6x (ranging from 24.0x to 56.6x). These reads were used to *de novo* assemble 46 partially-phased haploid genomes. This marks the first time *de novo* assemblies of Pacific ancestry genomes have been created, resulting in highly accurate and contiguous assemblies with an average QV of 55.0, or approximately one error per 300,000 base pairs. The assemblies had an average N50 of 40.7 Mb (ranging from 14.2 Mb to 83.0 Mb), with many assemblies nearing the GRCh38 N50 value of 58 Mb. The contiguity of these assemblies is comparable to those constructed by the HPRC, which had an average NG50 of 40 Mb^6^, and the Chinese pangenome assemblies with an average N50 of 36 Mb^12^. These assemblies are nearly complete, with an average length of 2.98 Gb (95.5% of the T2T-CHM13 reference length).

We identified protein coding gene duplications specific to the 46 Pacific ancestry assemblies. One of these genes, *NPHP1,* was observed in 3 individuals and is known to be associated with renal disease^26^. Additionally, we found a lower frequency of *CLPS* duplications in the Pacific ancestry assemblies compared to the HPRC assemblies, which encodes pancreatic colipase and has been linked to T2D^27^. These duplications represent genetic variation with disparate distributions across populations and could potentially contribute to the increased rates of renal disease and T2D observed in individuals with Pacific ancestry.

From the 46 Pacific ancestry assemblies, we developed a pangenome reference that contributed an additional 30.6 Mb of novel sequence absent from the T2T-CHM13, GRCh38, and HPRC references. Of this novel sequence, 9.6 Mb was identified in two or more individuals with Pacific ancestry, indicating Pacific ancestry-specific polymorphic sequences. In total, we discovered 3.4 million small variants and 4,721 structural variants unique to this population. Mapping short-read sequence data to this pangenome reduced variant call errors and improved the recall of true variants for both SNVs and indels across the genome, especially in challenging-to-map regions. These improvements are consistent with those observed when using other population-specific pangenome references^11,12^

While creating this pangenome, some limitations should be acknowledged. The demographics of the pangenome, which include a majority of participants with Tongan and Samoan ancestry, reflect the Pacific ancestry of both the Utah Pacific Islander Diabetes Study and the population of Utah (69% of Utah’s Native Hawaiian and Pacific Islander community identify as either Tongan (39%) and Samoan (30%), while 7% identify as Native Hawaiians), but may not represent global Pacific ancestry demographics.

The creation of this pangenome reference was enabled by recent advances in long-read sequencing technology and bioinformatics tools, enabling highly accurate and contiguous genome assemblies with far less effort and resources than previously required. Although these tools are highly effective, they are not without imperfections, and without extensive manual curation, errors are likely to remain in these assemblies. However, state-of-the-art assembly analysis tools like CRAQ^10^ and Inspector^7^ offer reliable quality estimates for these assemblies and effectively pinpoint error-prone regions that can be excluded from downstream analyses.

This study aimed to deepen our understanding of genetic variation in individuals with Pacific ancestry by creating a pangenome resource tailored to this population. This pangenome reference significantly enhances variant calling accuracy in short-read sequence data. Further, it addresses gaps left by previous genome resources, offering a more precise tool for identifying previously inaccessible variants. Given the high prevalence of metabolic diseases among individuals with Pacific ancestry and the existing lack of comprehensive genetic data, our pangenome provides a crucial foundation for future research. It will enable more accurate genetic studies and potentially improve health outcomes through more informed precision medicine approaches.

## Methods

### Ethics and inclusion

Throughout the design and execution of this study, we actively engaged with local community leaders of Pacific ancestry to ensure the study was conducted in an ethical manner. This engagement included consultations with physicians and researchers, including coauthors of this study, who are familiar with the specific nature of this research, as well as the collection and storage of DNA sequencing data. All participants provided informed written consent prior to their involvement in the study. The consent process ensured that participants were fully aware of the study’s aims, procedures, and any potential risks and benefits. This study was reviewed and approved by the University of Utah Institutional Review Board (IRB_00159221), ensuring compliance with ethical standards and regulatory requirements. Additionally, we committed to transparency and reciprocity by sharing the study findings with the community and acknowledging their invaluable contributions.

### Sample selection

Samples selected for sequencing were collected from individuals living in Utah with Pacific ancestry as part of the Utah Pacific Islander Diabetes Study (UPIDS). During sample collection, participants completed a questionnaire to identify the Pacific Islands of origin for each of their four grandparents. We selected 23 samples from UPIDS for long-read HiFi sequencing whose four grandparents originated from the same Pacific Island region. Additionally, the proportion of individuals included from each Pacific Island region was chosen to maximize diversity while still reflecting the Pacific ancestry demographics of Utah, with the majority of participants having Tongan and Samoan ancestry.

### DNA extraction and library prep for whole-genome sequencing

#### PacBio long-read sequencing

Blood was collected from participants and stored in plastic K_2_EDTA blood collection tubes (BD Vacutainer). The samples were then centrifuged, and the buffy coat was transferred to another K_2_EDTA blood collection tube, which was stored at -20°C until DNA extraction. Prior to extraction, samples were thawed, RBC lysis buffer (eBioscience) was added, and the remaining cells were pelleted. High molecular weight DNA was extracted and purified using the Qiagen Genomic-tip kit and protocol. The column-cleaned DNA was precipitated by adding 2x ethanol to the column eluate, mixing, and storing in a -20°C freezer overnight, followed by pelleting the DNA via centrifugation. After removing the ethanol, the DNA was dissolved in Low EDTA TE buffer and sheared on a Diagenode Megaruptor-3 to an average size of 15-17 kb. Fragment size was verified on an Applied Biosystems Femtopulse. The DNA was further isolated using 0.5X AMPure and prepared for sequencing using the Procedure-checklist-Preparing-whole-genome-and-metagenome-libraries-using-SMRTbell-prep-kit-3.0 protocol. Sequencing was performed on the Revio 3 via the SMRTLink (v12.0) software. Samples were sequenced to a minimum HiFi yield of 86 Gb (∼30x genome coverage). Any samples with less than 86 Gb were re-sequenced, and data from both runs were combined to achieve the minimum required yield. The average genome sequencing depth of these reads aligned to T2T-CHM13 was 32.57X.

#### Illumina paired short read sequencing

Genomic DNA from the same 23 blood samples selected for long-read sequencing was extracted using a standard phenol-chloroform DNA extraction protocol. Sequencing was performed at the Broad Institute’s Clinical Laboratories using PCR-free library construction and paired-end (2×150 bp) sequencing on the Illumina NovaSeq X Sequencing System. Aligning the resulting reads to GRCh38 with Illumina’s DRAGEN software revealed an average genome sequencing depth of 32.74X.

### *De Novo* genome assembly and correction

Unaligned BAM files of HiFi long reads were first converted to FASTQ files using Bedtools bam2fastq (v3.1.1)^31^. For samples that were re-sequenced, the resulting FASTQ files were merged into a single file prior to genome assembly. We used a total of 23 FASTQ files, filtering out adaptor-contaminated reads with HiFiAdapterFilt (v3.0.1)^32^ software using default parameters. Next, we utilized Hifiasm (v0.19.6)^8^ to *de novo* assemble the initial 46 partially-phased haploid assemblies with the following command: "*hifiasm -o $assembly $input.reads -t 64*". The quality of these assemblies was evaluated using the genome assembly analysis tool Inspector(v1.0.1)^7^. Inspector uses a reference-free approach by aligning HiFi long read sequences to their respective assemblies and identifying inconsistencies. It also performs reference-based analyses by aligning the assemblies to a linear reference and reporting genome coverage statistics. The following command was run twice, once using GRCh38 and once using T2T-CHM13 as the reference: “*inspector.py -c $input.fasta_file -r $input.fastq$ -o $output_dir$ --datatype hifi --thread 64 --ref $reference_genome”.* The output summary files for each assembly provided data on N50, total base pair length, read-to-assembly mapping rates, quality values (QV), number of contigs, number of errors (structural and small), and coverage statistics for both GRCh38 and T2T-CHM13. After evaluation, assemblies were corrected using Inspector. Inspector corrects the assemblies by using read-to-assembly alignments to identify substituted, collapsed, and expanded regions based on read depth. Regions with structural errors were deconstructed and reassembled using local reads and the Flye genome assembler^33^. This was achieved with the following command: “*inspector-correct.py -i $input_dir --datatype pacbio-hifi -o $output_dir --thread 64*”.

### Evaluating assembly quality

After Inspector correction, assemblies were again evaluated by Inspector for quality using the Inspector evaluation script described previously. To avoid the potential bias of using Inspector to re-evaluate the same assemblies it corrected, assemblies were also evaluated post-correction using CRAQ (v1.0.9)^10^. CRAQ aligns both Illumina short-read and PacBio HiFi long read data to their respective assemblies and identifies structural and regional errors based on groupings of clipped reads in the same genomic regions. Regional errors are interpreted as smaller local errors, while structural errors represent larger assembly misjoins. This approach aims to mitigate the impact of error-enriched regions on the overall assembly quality score. Structural and regional assembly quality indicators (AQI) were calculated using the following command: "*craq --genome $asm -sms $long-reads -ngs $short_reads --sms_coverage 30 --ngs_coverage 30 -- break T --map map-hifi --mapq 20 --output_dir $out --thread 64* ".

Compleasm (v0.2.6)^23^ was used to analyze the completeness of assemblies, by mapping universal single-copy orthologs for primates to each assembly downloaded from the following location: “https://busco-data.ezlab.org/v5/data/lineages/”. This was performed using default parameters and the following command: “*singularity exec docker://huangnengcsu/compleasm: v0.2.6 compleasm run -a $asm -o $name -l primates -t 32*”

### Characterizing unaligned assembly sequences

Assembly contigs were aligned to GRCh38 and T2T-CHM13 through Inspector (v1.0.1), which uses minimap2^34^ with preset parameters "-x asm5". Unaligned portions of each contig were extracted from the resulting alignment (BAM) files, and the number of base pairs was summed to calculate the total length of unaligned sequence. Further, the unaligned sequences of each BAM file were converted into FASTA format and analyzed by RepeatMasker (v4.1.6)^24^ using NCBI/RMBLAST (v2.14.0) search engine and the Dfam database (v3.8)^35^ to determine the portion of these sequences that are repetitive via this command: "*RepeatMasker -e rmblast --dir "$out_dir" "$seqfile*".

### Copy number variant analysis

To identify protein-coding gene duplications in our 46 assemblies, we utilized Liftoff (v1.6.3)^25^ to annotate genes with at least 90% matching sequence identity. We executed the following command: "*liftoff -p 64 -sc 0.90 -copies -db $gtf -o output.gtf -polish $asm $ref*". The GTF file used to create the database was obtained from Gencode and included comprehensive gene annotations for GRCh38 reference chromosomes only. Whole-gene duplications were identified by selecting genes in the GTF output that had a value greater than 0 in the "extra_copy_number" field. To determine whole-gene duplications specific to our 46 Pacific ancestry assemblies, we conducted the same analysis on the 94 HPRC assemblies and compared the results with our Pacific ancestry assemblies. In this study, gene duplication "frequency" refers to the number of assemblies with at least one duplicated copy of the same gene, divided by the total number of assemblies in its respective dataset.

### Pangenome graph construction

The pangenomes were created using the Minigraph-Cactus pipeline^28^ (Cactus v2.7.2). Minigraph-Cactus generates assembly-based pangenome graphs by aligning de novo assemblies to a reference. The process begins by mapping assembly contigs using Minigraph to create a structural-variation-only graph. The assemblies are then remapped to this graph to incorporate single nucleotide variations, while sequences longer than 10 kb that remain unaligned are excluded to remove highly repetitive regions, such as those in telomeres and centromeres. All graphs were based on the T2T-CHM13 reference but also included a version of GRCh38 without alternate contigs as a "reference-sense" path in the graph. We utilized this pipeline to create all pangenome references analyzed in this study using the following command: "*cactus-pangenome./temp $seq_file --outDir $out_dir --outName $out_name --reference CHM13 GRCh38 -- mgMemory 250Gi --consMemory 250Gi --indexMemory 250Gi --haplo clip --gbz --gfa --giraffe filter clip --vcf --odgi --chrom-og --viz --draw –restart*". The Pacific ancestry pangenome references were created using the 46 Pacific ancestry assemblies corrected by Inspector. The HPRC graph used in this study was constructed using the 94 HPRC Year 1 v2 assemblies published by the HPRC^6^. The combined Pacific ancestry + HPRC graph was constructed using the 46 Pacific ancestry and the 94 HPRC assemblies together in one graph.

The output of the Minigraph-Cactus pipeline includes graphs with variation included from every assembly, including singleton sequences. In addition, we generated allele-frequency-filtered graphs that retain only the nodes (sequences) traversed by at least 2 to 14 haplotype paths. These filtered graphs are less complex and provide better performance for read alignment using Giraffe^9^. We created these allele-frequency-filtered graphs for each of the three graphs (Pacific ancestry, HPRC, and Pacific ancestry + HPRC) using the following command "*cactus-graphmap-join./$freq.temp --filter $freq --gfa filter --vg $vg_dir/*.vg --outDir $out_dir -- outName $out_name --reference CHM13 GRCh38*". The complexity of these graphs was calculated using vg stats^36^.

### Pangenome cumulative growth curve

The pangenome growth curves and the number of novel base pairs and nodes added with each sample were calculated using Panacus (v0.2.3)^37^. The default graphs produced by the Minigraph-Cactus pipeline were used as input for this analysis using the following command: "*panacus ordered-histgrowth --count {bp/node} --order $ordered_list --exclude corrected.paths.CHM13.txt --subset corrected.paths.haplotypes.txt --output-format table -- coverage $coverage --groupby-sample --threads 64 $gfa > $gfa.ordered-histgrowth.bp.tsv*".

Coverage thresholds of 1, 2, and 22 were selected to represent singleton, common (5%), and core (95%) sequences, respectively. In creating the ordered list of Pacific ancestry samples, we grouped individuals from the same Pacific region together and prioritized regions with the greatest number of samples. The same analysis was performed for the Pacific ancestry + HPRC graph, and the ordered list included the HPRC samples first, followed by the Pacific ancestry samples. Coverage thresholds differed for this graph, including 1, 2, 4, and 67, which represented singleton, polymorphic (allele count >1), common (5%), and core (95%) sequence, respectively.

### Characterizing variation of the pangenome graphs

To identify the variation unique to individuals with Pacific ancestry, we utilized the Pacific ancestry + HPRC pangenome graph and the VCF created using vg deconstruct and vcfbub via the" --vcf" option of Minigraph-Cactus pipeline against the T2T-CHM13 reference. We then decomposed and normalized this VCF using BCFtools (v1.16)^38^. The VCF was subsequently split into two separate files: one containing variants observed exclusively in Pacific ancestry samples and another containing variants specific to HPRC samples. We then identified variants unique to each VCF as well as those shared, using the intersection functionality of BCFtools: "*bcftools isec -c none -p unique_subset -Oz subset.vcf.gz control.vcf.gz*". Variants were then stratified into small and structural variants based on size (structural variants >50bp).

### Aligning short read sequences to the pangenome references and calling variants

Illumina short reads were aligned to the pangenome references using the personalized pangenome approach of Giraffe^39^ (vg v1.55.0), which relies on haplotype sampling. First, paired FASTQ files of Illumina short read sequences were sampled for k-mers using the following command: "*kmc -k29 -m128 -okff -hp -t64 @./fq_list.txt $name."* The resulting KFF files were used to sample the haplotypes of the pangenome graphs and create personalized pangenome graphs for each sample. To create the personalized pangenome graphs and map short reads to them, we used the following command: "*vg giraffe -Z $gbz -f $fq1 -f $fq2 -o bam –sample $sample --progress --kff-name $sample.kff --haplotype-name $hapl.hapl -R ’ID:1 LB:lib1 SM:${sample} PL:illumina PU:unit1’ --ref-paths $paths > $sample.$name.bam*”. In addition to aligning short reads to the pangenome references, we also aligned reads to the T2T-CHM13 linear reference for downstream variant benchmarking. This was achieved using BWA-MEM (v 0.7.17)^40^ and the following command: *bwa mem -t 64 hs1.fa -T 0 "${sample}_R1.fq.gz" "${sample}_ R2.fq.gz" > "${sample}_T2T.sam*". The individual alignment files for each reference were examined for read mapping quality using Samtools (v1.17)^38^.

Variant calling was performed using DeepVariant (v1.6.0)^41^. BAMs produced from Giraffe alignment were used as input, and individual VCF files and genome VCF files (gVCF) were produced using the following command: *singularity exec -H $(pwd) docker://google/deepvariant:1.6.0 /opt/deepvariant/bin/run_deepvariant --model_type=WGS -- ref=$ref --reads=$bam --output_vcf="$bam_name.vcf.gz" --output_gvcf="$bam_name.gvcf.gz" --make_examples_extra_args="min_mapping_quality=1,keep_legacy_allele_counter_behavior =true,normalize_reads=true".* VCF files were then filtered to include only "PASS" variants before use in downstream applications.

### Benchmarking small variant calls

Pseudo-truth variant call sets were created using Dipcall (v0.3)^29^ to call variants from assemblies aligned to the T2T-CHM13 reference. The five most contiguous Pacific ancestry samples were selected for benchmarking based on their combined N50 values from both haploid assemblies: UP001, UP003, UP004, UP009, and UP022. Assemblies were aligned to the T2T-CHM13 reference and variants were called using the following commands: */uufs/chpc.utah.edu/common/home/u0962117/software/dipcall.kit/run-dipcall $sample $ref $hap1 $hap2 > $sample.mak make -j2 -f $sample.mak*" In addition to calling variants, Dipcall identified confident regions in the VCF by selecting base pairs covered by at least one alignment of ≥ 50 kb with a mapping quality score > 5 from each parent, as long as these areas were not covered by other alignments of ≥ 10 kb^42^. For downstream benchmarking with Hap.py, variant calls were restricted to these confident regions, and variants larger than 50bp were removed.

We then analyzed concordance between the variant calls of the Dipcall truth sets and the variant calls from the short read data of the same samples aligned to T2T-CHM13, the Pacific ancestry reference, and the HPRC reference using Hap.py (v0.3.15)^43^: "*python2.7 $(which hap.py) $truth $query -f $dip_bed --threads 64 -r $ref --write-vcf --pass-only --engine xcmp --stratification $tsv --roc QUAL --preprocess-truth -o happy.$name*". The analysis produced concordance metrics (including recall, precision, and F1 scores) for indels and SNVs separately, as well as the number of true positives, false positives, and false negatives. Concordance was further stratified into five genomic regions using BED files produced by the Genome in a Bottle (GIAB) consortium (v3.4) for the T2T-CHM13 reference:

1) CHM13_alldifficultregions.bed.gz (“Difficult regions”)
2) CHM13_alllowmapandsegdupregions.bed.gz (“Low map and seg dup”)
3) CHM13_notinalldifficultregions.bed.gz (“Easy regions”)
4) CHM13_notinalllowmapandsegdupregions.bed.gz (“Not in low map and seg dup”)
5) CHM13_refseq_cds.bed.gz (“Coding Sequence”).

### Principal component analysis (PCA)

To identify the variation unique to individuals with Pacific ancestry, we utilized the Pacific ancestry + HPRC pangenome graph and the VCF created using vg deconstruct and vcfbub via the" --vcf" option of Minigraph-Cactus pipeline against the T2T-CHM13 reference. GRCh38 variants were first removed from the VCF, and the remaining variants were filtered to include only biallelic SNVs with less than 10% missingness. This VCF was then filtered using Plink(v1.90)^44^ to remove variants with an allele frequency < 0.05 and a significant deviation from expected allele frequencies (p-value < 0.00000001) using the following command: *"plink -- noweb --allow-no-sex --bfile $DATA --maf 0.05 --hwe 0.00000001 --make-bed –out ${OUT}/CommonAlleles".* Linkage disequilibrium pruning was performed using this command: "*plink --noweb --allow-no-sex --bfile ${OUT}/CommonAlleles --indep-pairwise 50 5 0.5 -out ${OUT}/prunedsnplist*". The resulting list of SNPs was used as input to compute principal components via Plink: "*plink --noweb --allow-no-sex --bfile ${OUT}/CommonAlleles --pca -- extract ${OUT}/prunedsnplist.prune.in --make-bed --out ${OUT}/PCA*".

## Data availability

This manuscript represents only a draft of the full study to be released in the future. The Pacific ancestry pangenome and associated assemblies will be made available in the final versions of this study.

## Acknowledgments

The authors would like to acknowledge grant support from the National Institutes of Health (DK128641 to MGP, DK139787 to WHL, KLR, and MGP, MD016482 to MGP, and 5T32DK110966 to ML), the National Kidney Foundation of Utah & Idaho (to MGP), the Center for Genomic Medicine at the University of Utah Health (to MGP), the Ben B. and Iris M. Margolis Foundation (to MGP), and the University of Utah Diabetes and Metabolism Center (to CL). We also acknowledge the support of our community partners in Utah and the valuable contributions of all participants of the Utah Pacific Islander Diabetes Study.

## Author contributions

CL, KLR, WLH, and MGP designed the study; CL, KLR, WLH, and MGP wrote the manuscript; CL, JML-G, DS, ML, E-MCW, CDCD, JPT, KLR, WLH, and MGP analyzed the data and generated all the Figures and Tables; MGP, MHP, EJT, JT and KL provided DNA samples and clinical phenotype data; CL, JML-G, DS, ML, PGR, JPT, KLR, WLH, and MGP assisted in the data interpretation. All authors edited and have read and approved the manuscript.

