## Supplementary Figures for "A Draft Pacific Ancestry Pangenome Reference"

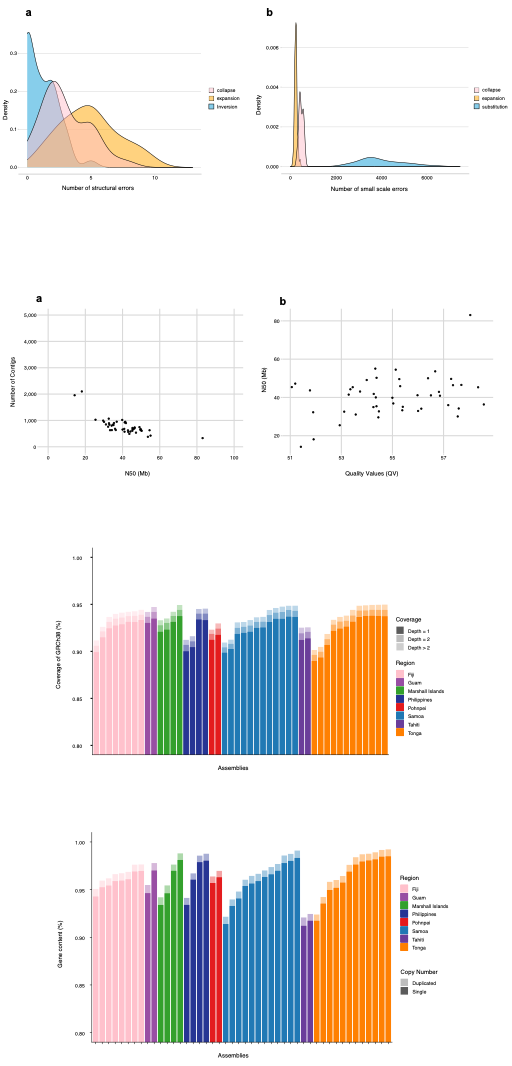

**Supplementary Figure 1. Coverage of GRCh38 by the 46 Pacific ancestry assemblies**

The Y-axis shows the percentage of genes present in each assembly determined by compleasm, for each assembly on the X-axis. The color intensity indicates the copy number of the genes identified in the assemblies. The highest color intensity represents genes observed only a single time, and the weaker intensity represents genes duplicated at least once. The average percentage of total gene content in all assemblies is 96.6% (including single and duplicated, and excluding fragmented and missing).

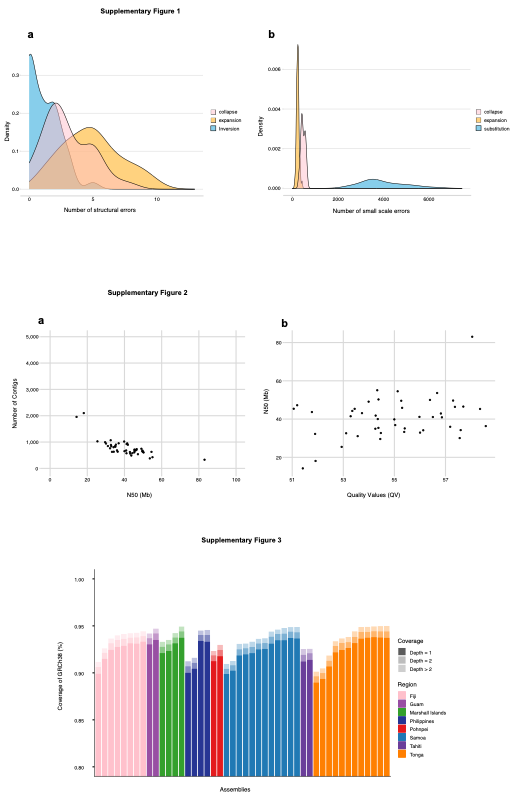
**Supplementary Figure 2. Small and structural errors in the 46 Pacific ancestry assemblies**

**a)** Density of small-scale errors identified by Inspector post-correction, categorized into collapsed regions (pink), expanded regions (yellow), and base pair substitutions (blue).

**b)** Density of larger structural errors also identified by Inspector post-correction, categorized into collapsed regions (pink), expanded regions (yellow), and inversions (blue).

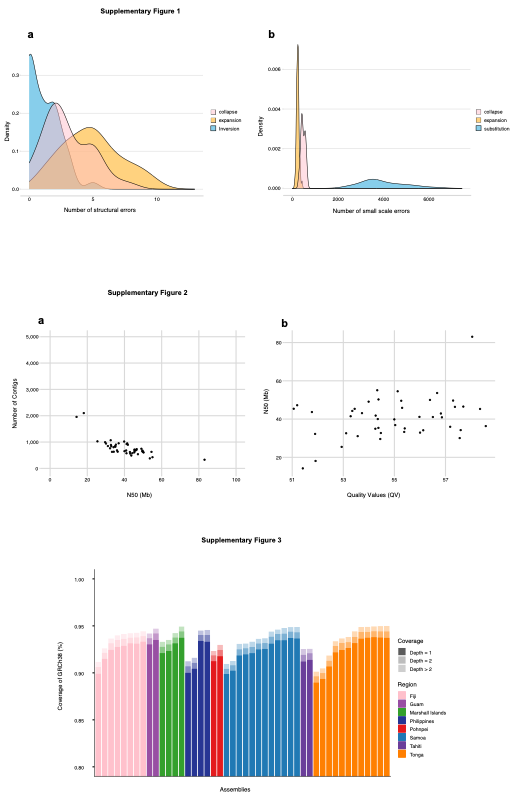

**Supplementary Figure 3. Quality and Contiguity of the 46 Pacific ancestry assemblies**

**a)** Plot of the number of contigs per assembly on the *Y* axis and the contiguity measured by N50 (the length of the shortest contig at 50% of the total assembly length) on the *X* axis. Average number of contigs for all 46 assemblies is 782, and average N50 is 40.7Mb.

**b)** Plot of N50 values on the *Y* axis and quality values (QV) identified post-correction by Inspector on the x axis. Average QV score is 55.0.

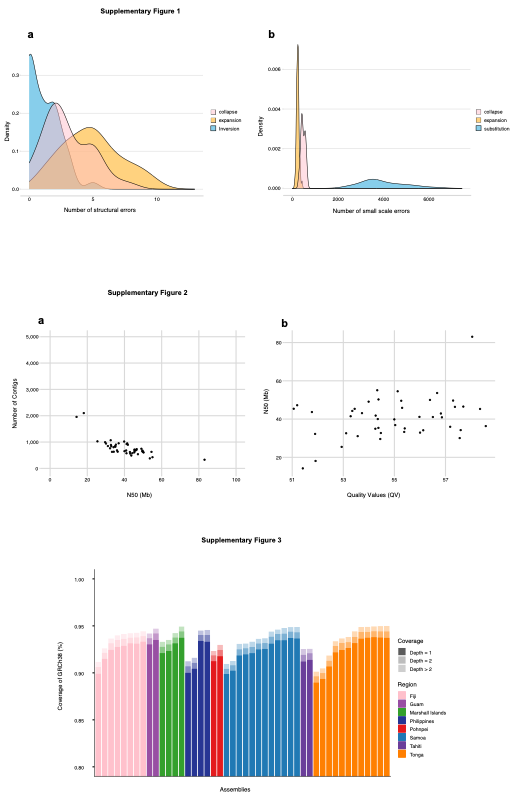

**Supplementary Figure 4. Coverage of GRCh38 by the 46 Pacific ancestry assemblies**

The Y-axis shows the percentage of GRCh38 covered by each Pacific ancestry assembly, with assemblies ordered by ancestry on the X-axis. The color intensity indicates the percentage of the genome covered by one portion of the assembly (highest intensity), two portions of the assembly (medium intensity), or more than two portions of the assembly (lowest intensity). The average coverage for GRCh38 was 93.5%.

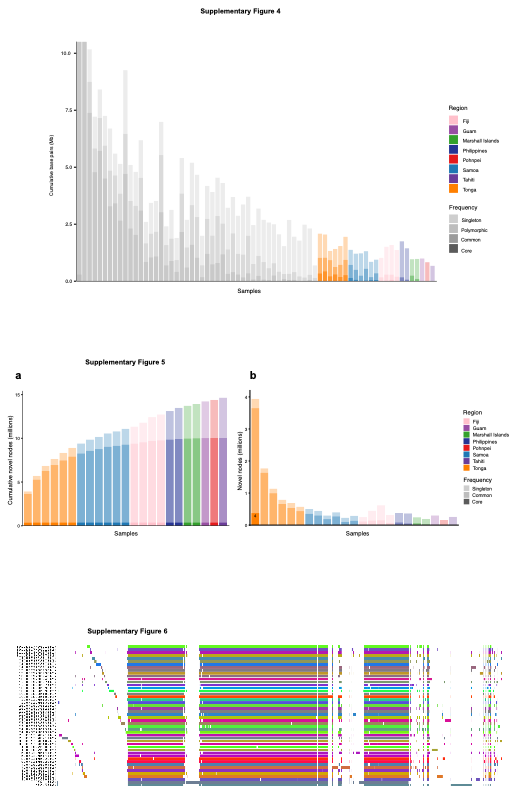

**Supplementary Figure 5. Novel base pairs contributed by each individual assembly to the combined HPRC + Pacific ancestry pangenome variation graph.**

The Y-axis shows the number of novel base pairs added to the combined HPRC + Pacific ancestry pangenome per sample, ordered from left to right. HPRC samples are in gray, while Pacific ancestry samples are colored. The color intensity indicates the frequency of the novel sequence: singleton sequence (observed only once), polymorphic sequence (observed more than once), common sequence (observed in at least 5% of samples), and core sequence (observed in at least 95% of samples).

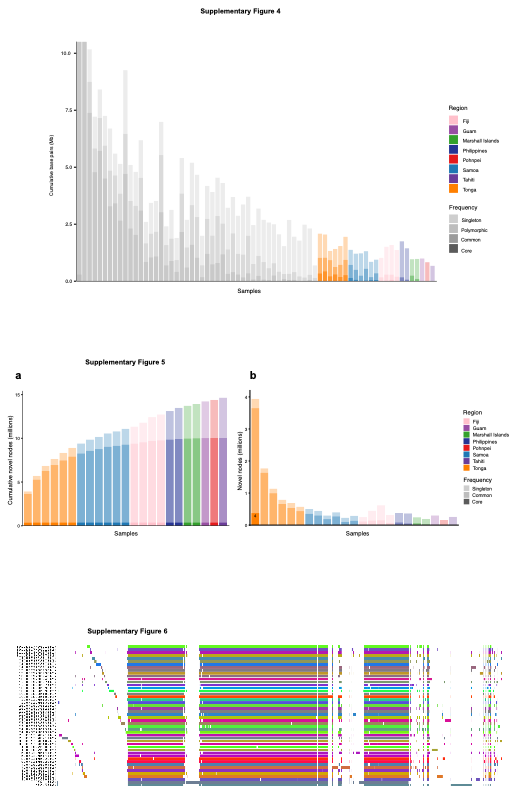

**Supplementary Figure 6. Novel sequences added to the T2T-CHM13 and GRCh38 references in the Pacific ancestry pangenome graph.**

**a)** The Y-axis shows the cumulative number of novel nodes (sequences) added to the Pacific ancestry pangenome graph per sample, ordered by ancestry. The color intensity indicates the frequency of the novel sequence: singleton sequence (observed only once), common sequence (observed in at least 5% of samples), and core sequence (observed in at least 95% of samples).

**b)** Similar to figure a), except the Y-axis shows the number of novel nodes added for each individual sample rather than the cumulative number.

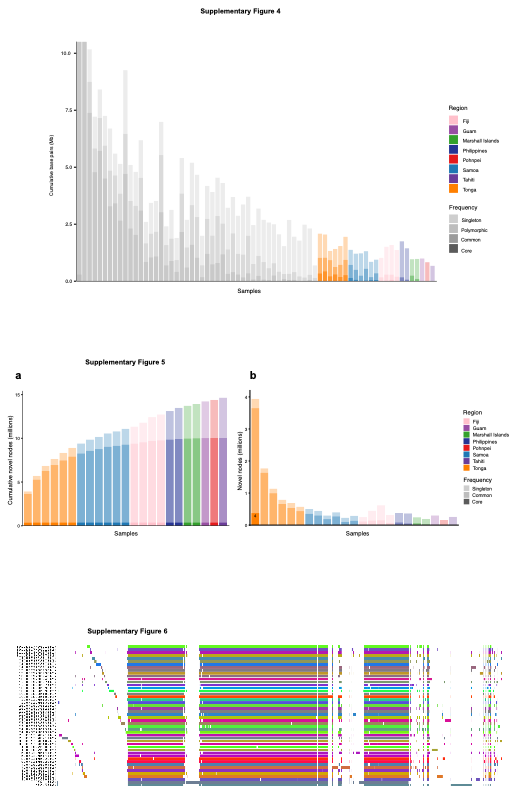

**Supplementary Figure 7. Visual representation chromosome 1 of the Pacific ancestry pangenome graph.**

This plot shows the alignment of sequences (colored lines) from each assembly, including the linear reference GRCh38, to the T2T-CHM13 reference within the Pacific ancestry pangenome graph. Alignments are displayed vertically in line with the T2T-CHM13 sequences. This graph was generated using the Minigraph-cactus pipeline and visualized with ODGI viz^1^.

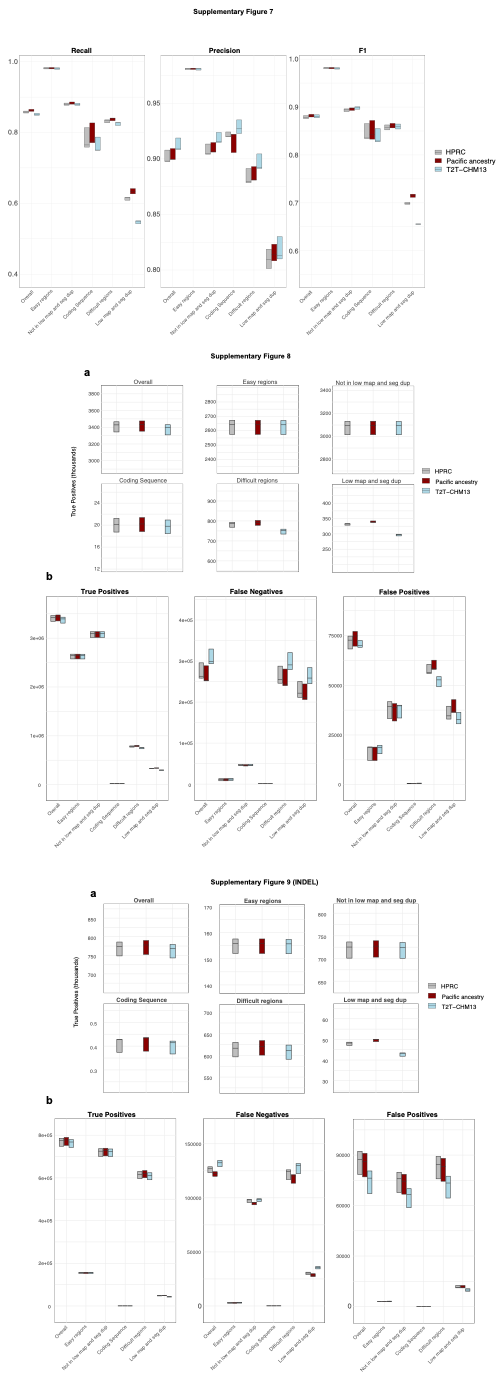

**Supplementary Figure 8. Evaluating improvements in indel calling by aligning whole-genome short-read sequences to the Pacific ancestry pangenome reference.**

Box plots showing recall, precision, and F1 scores for indels called from short-read sequence data of five Pacific ancestry samples, and stratified by GIAB-specified genomic regions^2^. ” Low map” refers to low mappability, and “seg dup” refers to segmental duplications. The samples are aligned to either the HPRC pangenome reference (gray), the Pacific ancestry pangenome reference (red), or T2T-CHM13 (blue).

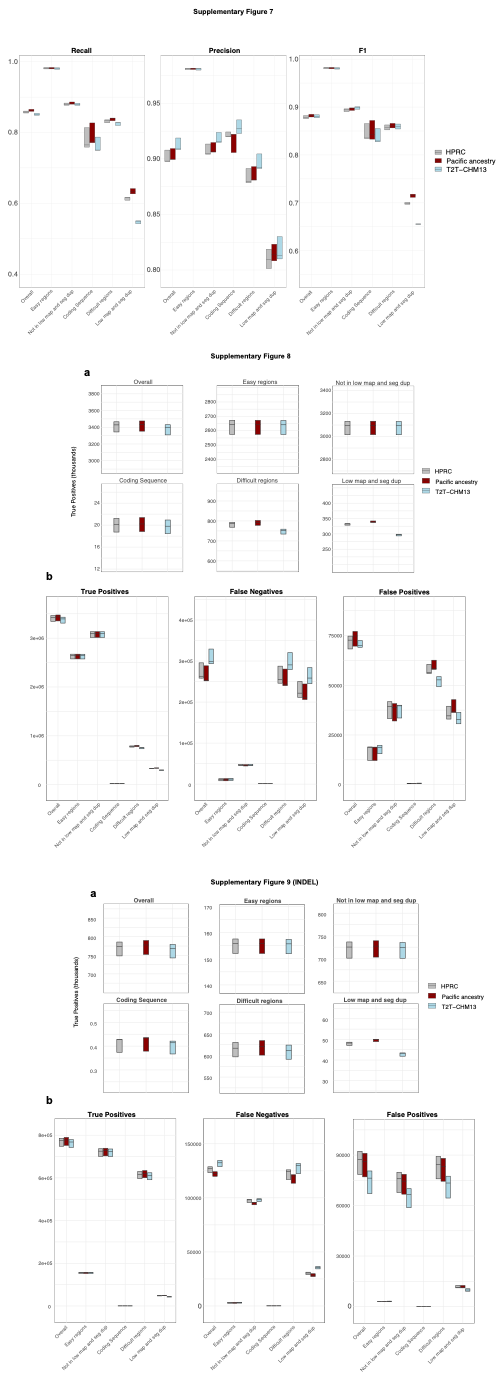

**Supplementary Figure 9. Improvements in SNV calling accuracy using short-read sequences aligned to the Pacific ancestry pangenome reference.**

**a)** Box plots showing the number of true positive SNV calls from short-read sequence data of five Pacific ancestry samples, stratified by GIAB-specified genomic regions^2^. ” Low map” refers to low mappability, and “seg dup” refers to segmental duplications. The samples are aligned to either the HPRC pangenome reference (gray), the Pacific ancestry pangenome reference (red), or T2T-CHM13 (blue).

**b)** Box plots showing the number of (from left to right) true positive, false negative, and false positive SNVs calls from the same dataset across different genomic regions.

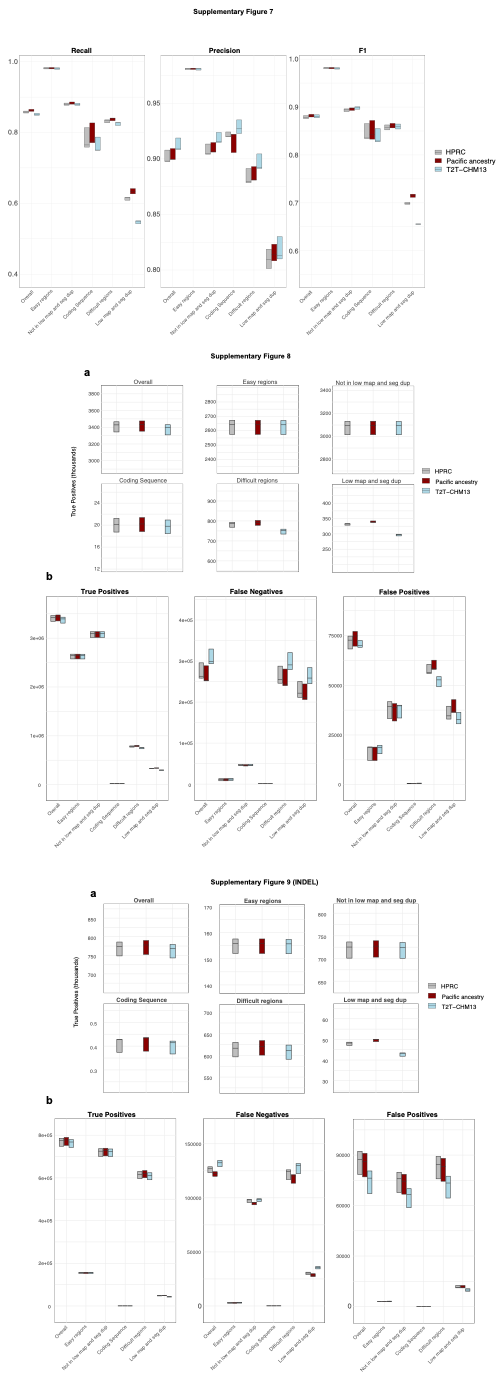

**Supplementary Figure 10. Improvements in indel calling accuracy using short-read sequences aligned to the Pacific ancestry pangenome reference.**

**a)** Box plots showing the number of true positive indel calls from short-read sequence data of five Pacific ancestry samples, stratified by GIAB-specified genomic regions^2^. ” Low map” refers to low mappability, and “seg dup” refers to segmental duplications. The samples are aligned to either the HPRC pangenome reference (gray), the Pacific ancestry pangenome reference (red), or T2T-CHM13 (blue).

**b)** Box plots showing the number of (from left to right) true positive, false negative, and false positive indel calls from the same dataset across different genomic regions.
